# Mechanism of molecular recognition revealed through dynamic drug binding pathways to SARS-CoV-2 main protease

**DOI:** 10.64898/2026.09.21.753329

**Authors:** Daniel Santos Perez, Grace Arhin, Yao Fu, Sean McCarty, Shu-Hang Lin, Terra Sztain

**Affiliations:** Department of Medicinal Chemistry, University of Michigan, Ann Arbor, Michigan 48104, United States; Biophysics Program, University of Michigan, Ann Arbor, Michigan 48109, United States

## Abstract

Characterization of drug-binding pathways remains experimentally limited by transient intermediates and computationally challenging due to long timescales intractable for conventional molecular dynamics. To address these challenges, we combined solution NMR titrations with weighted ensemble (WE) enhanced sampling simulations to resolve atomistic pathways of nirmatrelvir binding to the SARS-CoV-2 main protease. NMR titration revealed residue-dependent heterogeneity spanning fast, intermediate, and slow exchange regimes. WE simulations complement the NMR by providing insights into unassigned residues and adding time-resolved and three-dimensional structural context. We map key interactions along two distinct binding pathways, provide dynamic explanations for residues involved in resistance, and capture unique backbone conformations compared to those sampled in unbound or bound states. Our comprehensive binding model is consistent with a combined conformational selection and induced fit mechanism in which early transient contacts are made with residues E47 and L50 and allosteric motions are centered around residue V204 of the distal domain. This synergistic application of WE and titration NMR enables a more comprehensive characterization of drug binding than either method alone, providing an integrated framework that may have broader applicability to defining structure-kinetic relationships and guiding design of next-generation inhibitors.

## Introduction

The role of ligand binding kinetics is increasingly recognized as an important parameter for lead optimization, giving rise to the concept of structure-kinetic relationships (SKR), with particular emphasis on target residence time.^1^ More recently, the correlation between ligand association, k_on_, and drug efficacy has been established.^2,3^ Ecker and colleagues demonstrated that association kinetics can be rationally optimized through modulating the ligand desolvation barrier, representing a promising avenue for improving candidate drug efficacy.^4^ Due to the transient and dynamic nature of ligand association, this kinetic behavior remains difficult to characterize structurally, leaving it relatively underexplored.

The mechanism of ligand binding has historically been described by three models.^5^ First, the lock and key model portrayed the ligand and its receptor as static structures that complement each other’s shape perfectly.^6^ This has been largely superseded by the second, induced fit model, which describes how interactions with the ligand reshape the receptor inducing a well fitting bound conformation.^7^ Third, the conformational selection theory posits that protein conformations exist as an ensemble of states with different populations, and that a ligand can bind to a suitable pre-existing conformation and shift the equilibrium toward stabilizing that state.^8^ It is now widely accepted that proteins likely use a combination of the latter two models, where a productive encounter complex is first formed via conformational selection, which induces the conformational changes necessary for stable binding.^9–13^

Many structural techniques provide static snapshots that do not capture these dynamic phenomena. Solution nuclear magnetic resonance (NMR) spectroscopy uniquely enables high-resolution observation of molecules in a native-like, dynamic environment. One dimensional saturation transfer difference (STD)-NMR^14^ and two dimensional heteronuclear single quantum coherence (HSQC) titration are commonly used NMR experiments for studying ligand binding interactions. These can be used to determine whether a ligand binds, with what rates and affinities, and, in the case of HSQC, which protein residues are involved through chemical shift perturbation (CSP) mapping.^15^ In a standard ^1^H-^15^N HSQC spectrum, each cross peak typically corresponds to a specific residue in the protein backbone that is sensitive to the protein’s local chemical environment. The pattern of peak movement during an HSQC titration provides useful information about the exchange kinetics between unbound and bound states. Due to challenging peak assignment and overlapping peaks that frequently occur when sizes exceed a couple hundred residues, it is not feasible to depend exclusively on NMR for characterizing the structural complexities of ligand recognition. NMR and molecular dynamics (MD) simulations are frequently combined to construct a synergistic view of structure and dynamics in biological systems.^16–19^

Using MD to characterize protein conformational ensembles has been instrumental in drug discovery. A notable example is the discovery of raltegravir. Simulations of the HIV integrase identified a rare cryptic pocket conformation, which could be stabilized by a small molecule scaffold.^20–22^ This approach, leveraging conformational selection, is now widely used in computer-aided drug design.^23–25^ Induced fit conformations that are only accessible in the presence of an appropriate binding molecule have been captured using co-solvents or molecular fragments, which can also map hotspots on a protein surface.^26–28^ Capturing the complete dynamic mechanism of ligand association has been particularly challenging. Advances in enhanced sampling approaches have made it possible to accurately recover thermodynamic and kinetic properties in a feasible computational timeframe, such as Markov state models (MSMs),^29,30^ simulation enabled estimation of kinetic rates (SEEKR),^31^ ligand Gaussian accelerated molecular dynamics (LiGaMD),^32^ weighted ensemble (WE),^33^ and others,^34,35^ however there is a need to move beyond model systems to biomedically relevant targets, and for close coupling of computational predictions with experimental measurements.^36^

Here, we combine solution NMR titrations with WE enhanced sampling to study the association kinetics of the Paxlovid ingredient, nirmatrelvir, binding to the SARS-CoV-2 main protease. WE is well suited to preserving kinetic information because it does not modify the underlying force field, instead relying on splitting and merging of trajectories run in parallel with rigorous weight tracking. This enables direct estimation of rate constants from mean first passage times, with mechanistic resolution, at the level of the MD timestep rather than a lag time.

SARS-CoV-2 is the viral agent responsible for the COVID-19 pandemic. During the viral life cycle, the polyproteins pp1a and pp1b are produced encoding the non-structural components required for replication.^37,38^ Embedded within pp1a is the gene bearing the main protease (Mpro) also known as non-structural protein 5 (nsp5), or 3CLpro.^39^ Since Mpro has no human homologue,^40,41^ it has been the focus of extensive drug development efforts.^41–47^ Mpro shares ∼96% sequence identity with SARS-CoV and ∼50% with MERS-CoV main proteases, making it an attractive target for broad-spectrum inhibitor development.^48^ To date, Paxlovid^49^ is the only FDA-approved antiviral indicated for treatment of active SARS-CoV-2 infection that directly targets Mpro. Paxlovid is a combination therapy pairing the peptidomimetic inhibitor nirmatrelvir (NIR) with ritonavir, a CYP3A4 inhibitor which is included to slow metabolic degradation. Despite this success, resistance conferring mutations have been identified in clinical isolates from treated patients at hotspot residues such as E166, L50, and T21.^50^ Furthermore, ritonavir has several adverse drug interactions, preventing use in patients with certain conditions. The resistance potential and unfavorable pharmacokinetic properties indicate a need for next-generation and alternative Mpro inhibitors, motivating a deeper understanding of the mechanisms underlying Mpro inhibition.

Backbone resonance assignments for Mpro have been deposited to the Biological Magnetic Resonance Bank (BMRB)^47,51,52^ and subsequent HSQC studies have both identified novel Mpro-targeting ligands,^47,53^ and compared the conformational effects of NIR and prospective next-generation inhibitors using endpoint titration spectra.^54^ Here, we conduct an extensive HSQC titration series, resolving the exchange rate behavior for NIR binding to Mpro, and provide a three-dimensional context via complementation with molecular simulations of the binding pathway.

Several MD simulations have been carried out to investigate Mpro dynamics and the impact of substrate binding.^55–57^ Notably, WE simulations of peptide substrate binding carried out by Moritsugu et al. found dynamics of the flexible loop between residues 166-178 are key to substrate binding, highlighting residue E166 as a critical initial contact for association.^58^ The E166V mutation is one of the strongest NIR resistance mutations, though it also decreases Mpro activity and is often coupled with either the L50F or T21I compensatory fitness mutations.^59^ Wang et al. conducted LiGaMD simulations of NIR binding to Mpro, identifying key intermediate states along the binding pathway, including L50 and Q189 as important early contacts. They also calculated association and dissociation rate constants, resulting in a predicted K_D_ highly similar to experimentally reported values.^60^ LiGaMD builds on Gaussian accelerated MD (GaMD),^61^ which adds a harmonic boost potential to the potential energy surface to enhance sampling by decreasing barrier heights. In LiGaMD, thermodynamic reweighting is combined with Kramer’s rate theory, which accounts for the effects of diffusion on the transition rate.

Here, we apply the WE approach to study NIR association to Mpro with an unmodified force field, and interpret the sampled pathways in conjunction with NMR experiments to construct a comprehensive view of the dynamic binding process. We identify two primary NIR entry pathways which converge on a mechanism involving early interactions between the NIR trifluoro group and residue E47, and between the nitrile group and residue L50, followed by extensive NIR interactions with Q189 and engagement with residues S46, M49, M165 and/or E166 committing NIR to productive binding followed by reorientation within the active site. Furthermore, we identify overlapping and distinct intermediate conformations during the binding pathway compared to unbound or bound simulations, consistent with a combination of conformational selection and induced fit dynamics. Finally, we integrate results from WE and NMR to construct an integrated model of drug binding, explaining slow exchange dynamics of the distal residue V204 by correlated motions with active site residues. Overall, our findings advance our understanding of NIR binding to Mpro, provide new insights into design of next generation inhibitors for treating COVID-19, and set the stage for future efforts in characterizing structure-kinetic relationships by combining WE simulations and titration NMR.

## Results and Discussion

### NMR Suggests Heterogeneous Exchange Kinetics during Ligand Recognition

To study the dynamic interactions between Mpro and NIR, we first acquired a series of ^1^H-^15^N heteronuclear single quantum coherence (HSQC) spectra of ^15^N-labeled Mpro with increasing concentrations of NIR (**Figure 1A**). Backbone resonances were assigned using published chemical shifts from the biological magnetic resonance bank entry (BMRB) as references.^47,52^ The titration was carried out until the protein was saturated with ligand as demonstrated by minimal changes in the spectra upon addition of higher NIR concentrations, which was observed at a ratio of 1.25 NIR/Mpro molar equivalents. CSP mapping for each residue reveals multiple exchange regimes during binding. These include fast exchange, where increasing NIR concentration results in displacement of a single peak representing the average of the bound and unbound states; slow exchange, where increasing amounts of NIR result in the appearance of a second peak corresponding to the bound state, which increases in intensity as the unbound peak decreases in intensity corresponding to the population of that state (**Figure 1B**); and intermediate exchange, where increasing NIR concentration results in peak broadening and disappearance, followed by appearance of the bound state peak, where the intensities cannot directly be mapped to the state populations (**Figure 1C**).^15^

**Figure 1:**
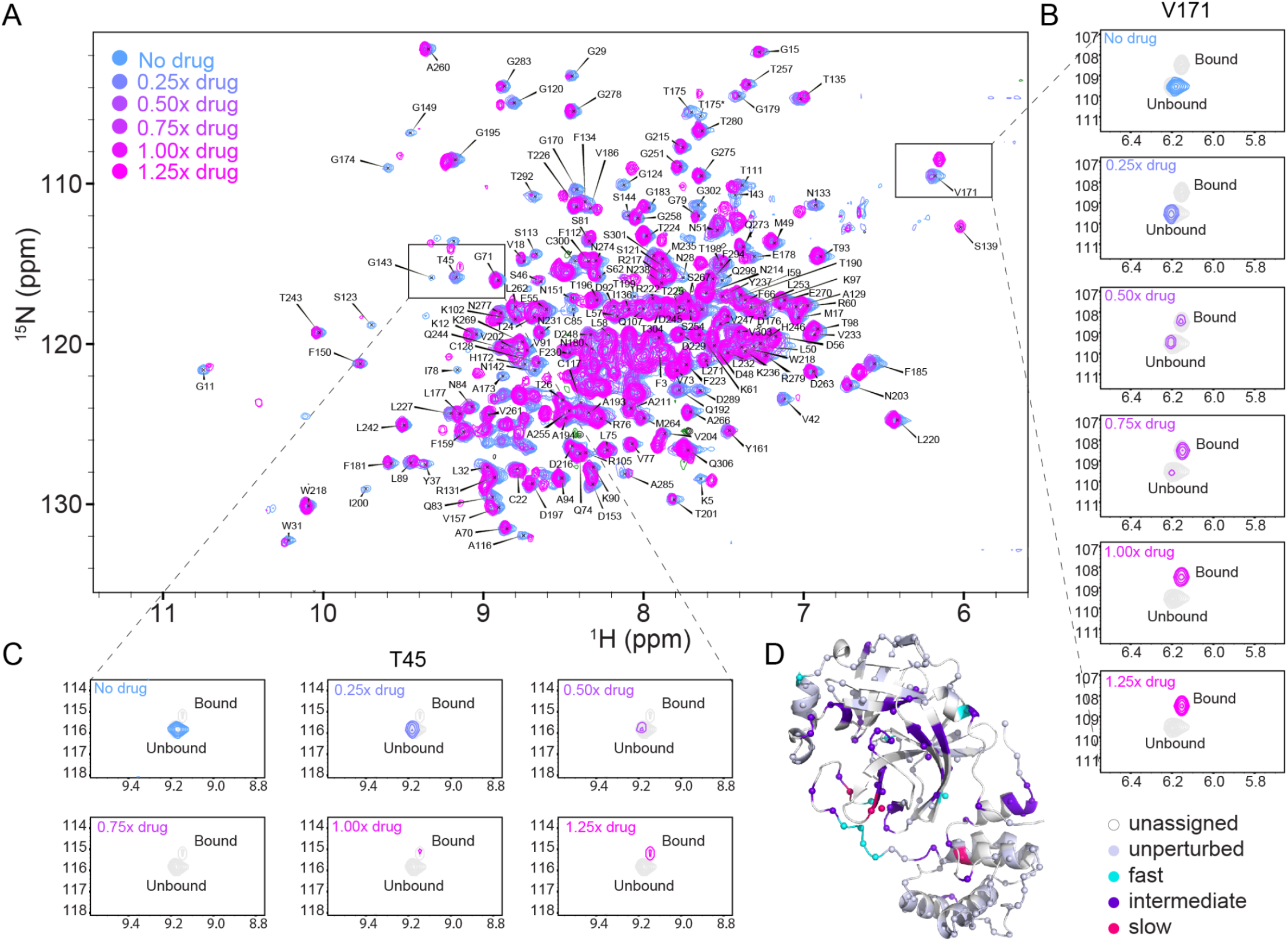
Mpro-WT recognition of NIR analyzed by NMR. (A) Overlaid ^1^H-^15^N HSQC spectra of wildtype Mpro at varying NIR concentrations. The asterisk on T175 shows an alternate conformation. (B) Expanded view of residue V171 to highlight slow exchange regime. Each titration point is colored relative to NIR concentration and remaining points are grayed out. (C) Expanded view of residue T45 to highlight intermediate exchange regime. Each titration point is colored relative to NIR concentration and remaining points are grayed out. (D) Perturbations are mapped onto the 3D Mpro structure. Residues observed in slow exchange are colored in hot pink, intermediate exchange in purple, fast exchange in cyan, unperturbed in slate, and unassigned in white.

Of the 156 assigned residues, 119 were unperturbed, 43 exhibited intermediate exchange kinetics, 9 displayed fast exchange behavior, and 5 were in slow exchange (**Table S1**). Perturbations span the active site and distal regions including the dimer interface, highlighting a complex mechanism of ligand recognition (**Figure 1D**). Perturbations far from the active site may be due to non-specific direct binding interactions, or they may be experiencing allosteric conformational rearrangement during binding.

Since NIR covalently inhibits Mpro by forming a bond with C145, we sought to determine whether the slow exchange behavior was due to increased affinity associated with the covalent bond formation. To test this, we repeated the NMR titration experiment with a C145A Mpro variant (**Figure S1**). The variant required 3 NIR/Mpro equivalents to reach saturation, compared to only 1.25 in the WT. Of the five residues that underwent slow exchange with the WT, only two, T135 and F185 remained in this regime. Residues G170, V171, and V204 exhibited intermediate exchange behavior. This suggests that the slow exchange behavior is not exclusively due to covalent bond formation. Additionally, several residues were perturbed via fast and intermediate exchange in the WT, yet unperturbed in C145A. Residues were also perturbed in fast or intermediate exchange in C145A that were not perturbed in WT (**Table S1**). These differences are non-localized and dispersed across the entire protein. From this information alone, it is not clear whether the observed CSP differences are due to differences in the conformational ensemble induced by the variant impacting residue kinetics and affinity, or whether they are due to preventing formation of a final bound conformation only possible when a covalent bond is formed. To further investigate the ligand recognition process at the atomistic level, we performed enhanced sampling simulations as described in the following section.

### Weighted Ensemble Simulations Illuminate Multiple Binding Pathways

The heterogeneity of exchange regimes observed by NMR spanning slow, intermediate, and fast exchange within the same spatial region suggests a kinetically diverse binding mechanism. To investigate these dynamics in greater detail, we used WE to generate an atomistic ensemble of NIR binding pathways, providing three-dimensional context to the observed two-dimensional NMR results. To generate a diverse ensemble independent of initial NIR configuration relative to Mpro, we constructed 64 independent systems with random initial NIR orientations 20 Å from the protein center of mass (**Figure S2**). Progress was monitored using a two-dimensional coordinate accounting for the distance between the nitrile warhead of NIR and the catalytic cysteine of Mpro, and the RMSD of NIR compared to the crystal structure of the covalently bound NIR. Since we were focused on characterizing the early association dynamics, a target state of below 5 Å was defined for both progress coordinates, and simulations were run under steady-state conditions, where trajectories that reach the target state are “recycled,” terminating the completed trajectory and initiating a new trajectory which inherits the weight of the terminated trajectory.^62^ Simulations were run until minimal fluctuations were observed in the calculated rate constant for several iterations. (**Figure S3**). The association rate constant stabilized around 1 x 10^4^ M^-1^s^-1^ after 103 iterations (**Figure S3**). Recently Chen et al. reported a K_on_ of 4.9 x 10^5^ M^-1^s^-1^ by globally fitting reaction-progress curves across a range of inhibitor concentrations to a mechanistic differential-equation model.^63^ This is highly similar to the K_on_ calculated from LiGaMD,^60^ 3.2 x 10^5^ M^-1^s^-1^, while our model is one order of magnitude slower. It is worth noting that estimations of K_on_ are highly sensitive to the definition of the bound state, and our 5 Å target state likely considered pathways complete during earlier encounter complex stages.

After 103 iterations, 7.1μs of total simulation time, 563 binding trajectories were generated. All successful trajectories originated from one of two initial NIR placements (**Figure S4**). The splitting events underlying WE simulation exploration result in a tree-like network of trajectory segments, which can each be traced to an initial state. While child trajectories that share a common ancestor are not strictly statistically independent, the stochastic nature of molecular dynamics ensures that trajectories diverge over time. Following sufficient simulation time after a splitting event, the microscopic configurations of sibling trajectories become uncorrelated, allowing them to be treated as independent samples of the binding pathway ensemble.^33,64^ To identify the predominant pathways from the ensemble based on their divergence, we performed hierarchical clustering, yielding 7 unique clusters. Trajectories within two clusters, 4 and 5, accounted for nearly 90% of the total recycled trajectory weight. Each cluster consists of trajectories originating from a different initial NIR placement so they contain no overlap in their evolutionary history. Accordingly, these were considered independent representative pathways which we analyze in greater detail in this section.

Pathways in cluster 5 originate from a large shared ancestry for the first 77 trajectories before splitting off, while cluster 4 begins to split after iteration 38, leading to a greater diversity of pathways (**Figure S5**). Visualization of all successful pathways reveals two distinct approach behaviors depending on the initial ligand placement (**Figure 2A**). The cluster 4 pathways do not directly approach the binding site, they start at a location where they must diffuse laterally across the length of the protein, subject to more of the protein’s forces and spending more time diffusing in solution before arriving at the appropriate conformation that leads to binding. Meanwhile, pathways in cluster 5 approach the binding site more directly. Snapshots from the most representative pathway from each cluster are shown in **Figure 2B**. In the cluster 4 pathway, NIR first engages the upper loop (residues 44–54) before finding its way deeper into the pocket towards the catalytic loop (162–174), and settling into position. In the cluster 5 pathway, NIR briefly engages with the upper loop before a long lasting interaction with the lower loop (184–194) until eventually finding its way into the pocket near the catalytic loop (**Figure 2C**).

**Figure 2:**
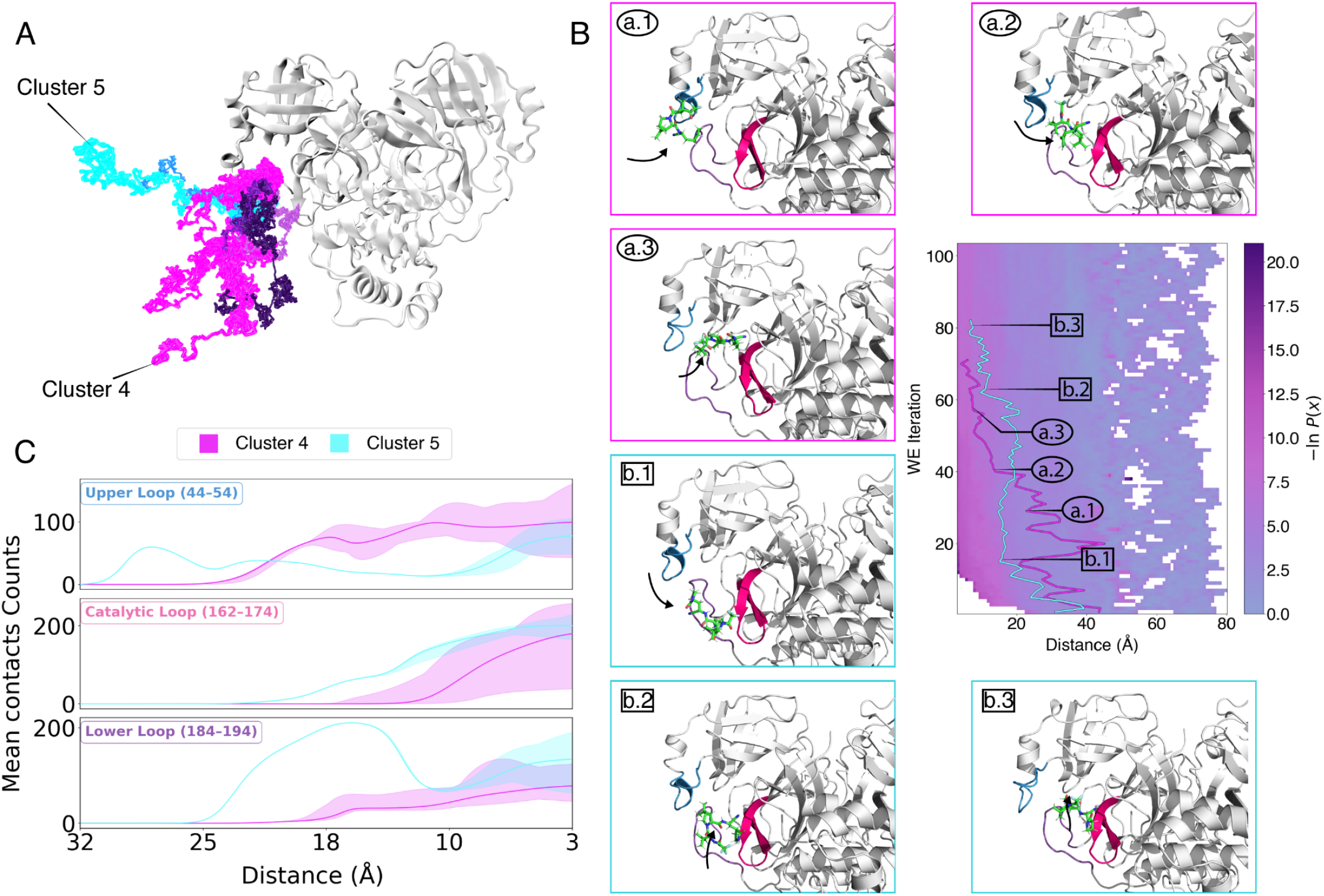
Overview of NIR-Mpro binding pathways from weighted ensemble simulations. (A) Overlay of the center of mass of ligand positions from all successful binding pathways. The seven colors correspond to each of the pathway clusters, with shades of blue or purple corresponding to distinct starting states. (B) Representative structural snapshots at key milestones along each pathway are shown in panels a.1–a.3 (magenta border, cluster 5) and b.1-b.3 (cyan border, cluster 4). Their positions along the pathway are indicated on probability distribution plotted as a function of distance of sulfur atom of residue C145 and the nitrile warhead nitrogen of NIR. For each representative structure, NIR is shown as green sticks, and the upper, lower, and catalytic loops are shown in blue, purple, and pink, respectively. (C) Mean contact counts of aggregated trajectory frames from each cluster as a function of distance of sulfur atom of residue C145 and the nitrile warhead nitrogen of NIR, shaded area show the minimum and maximum value.

We used ProLIF^65^ to quantify contacts between NIR and Mpro, identifying several residues contacted in over 90% of successful pathways in all clusters, those contacted in over 50% of successful pathways, and some cluster specific interactions (**Table 1**, **Table S2**). The majority of these residues line the binding pocket and are expected in the final bound conformation. However, residues H41, S46, E47, L50, N142, D187, and R188, are contacted in all pathways but are not detected as contacts in the bound crystal structure PDB 7SI9^66^ **(Figure S6)**. Instead, they are transiently contacted during the early stages of entry to the binding pocket. Conversely, all contacts identified in the static crystal structure were sampled during simulation.

**Table 1.** Residues contacted during MD simulation. Rows are separated based on whether they are present in the bound crystal structure, PDB 7SI9.

| Percentage of pathways and clusters in MD | Contacted in crystal structure? | Residues |
| --- | --- | --- |
| > 90%, both clusters | yes | M49, H164, M165, E166, P168, D187, Q189 |
|  | no | H41, S46, E47, L50, N142, D187, R188 |
| > 50%, both clusters | yes | G143, C145, H163, L167 |
|  | no | V186, Q192 |
| 100%, cluster 5 | yes |  |
|  | no | N51, V170, T190, A191 |
| > 50%, cluster 4 | yes |  |
|  | no | T25, T26, L27 |

These further away residues have been found to modulate Mpro catalytic activity, and susceptibility to NIR. Residue S46, for example, when mutated to S46F, has been shown to increase Mpro activity without compromising NIR sensitivity.^67^ Mutations in residue E47, such as E47N or E47K were reported to decrease or increase activity, respectively, while both minimally impacted NIR susceptibility.^67^ The L50F mutation increases catalytic efficiency without altering NIR response and is frequently observed as a compensatory mutation in variants that confer NIR resistance at a fitness cost.^59,68^ This suggests that the activity enhancing S46F, and E47K substitutions could potentially serve similar compensatory roles. A detailed description of the impact of mutations on Mpro activity and NIR susceptibility for each residue contacted in MD but not in the crystal structure can be found in the SI. Overall, this suggests that NIR susceptibility is shaped not only by residues in direct contact with the inhibitor in the covalent bound pose, but by a broader network of residues across the protein, and our simulations provide a plausible mechanistic basis for many residues previously found to impact Mpro activity and resistance.

### E47 and L50 may serve as early anchor points for nirmatrelvir binding

Next we characterized the residue-level interactions between NIR and Mpro for each pathway. For this analysis NIR was partitioned into chemical regions and motifs. These were defined as S1, containing the trifluoro group (S1-A), tert-butyl group (S1-B) and amide linker (S1-C); S2, the bicyclic proline motif; and S3, containing the amino nitrile group (S3-A) and the lactam group (S3-B) (**Figure 3A**).

**Figure 3:**
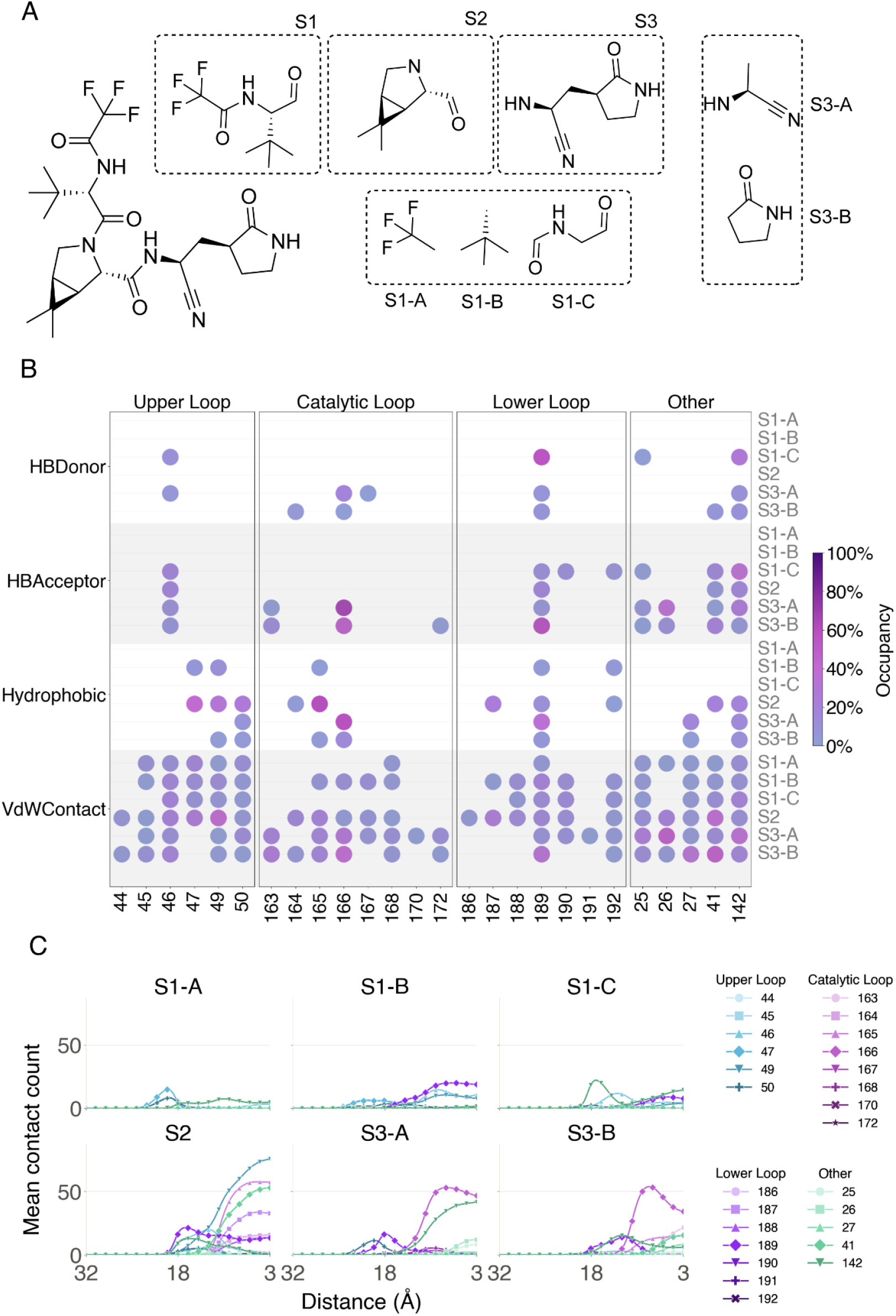
Fragment decomposition of NIR and per-fragment interaction fingerprints across Mpro loop regions for cluster 4. (A) Chemical structure of NIR (left) and its decomposition into three principal fragments used for the interaction analysis. (B) Contact-occupancy fingerprints between each ligand sub-fragment and individual Mpro residues, grouped by interaction type hydrogen-bond donor, hydrogen-bond acceptor, hydrophobic, van der Waals contact and by structural region, upper loop, catalytic loop, lower loop. Each point marks a detected interaction, color encodes occupancy measured as the number of frames a residue has a specific contact type out of all frames across the ensemble of successful pathways. (C) Mean contact counts of aggregated trajectory frames from each cluster as a function of distance of sulfur atom of residue C145 and the nitrile warhead nitrogen of NIR, each residue is represented by its own line and shape marker.

For pathways in cluster 4, NIR first engages with Mpro through transient hydrophobic interactions between the trifluoro group and upper loop residues E47 and L50. Next, the tert-butyl group interacts with E47 before migrating to residue Q189 of the lower loop and N142. Several other NIR regions interact with Q189 and N142 before settling into the bound conformation. Concurrently, upper loop residue S46 participates in both hydrogen bond (HB) donor and HB-acceptor contacts, engaging with several NIR regions simultaneously including S2 and S1-C, followed by S1-B. As binding progresses, these NIR regions interact with residue M49 in the upper loop, before interactions settle into the catalytic loop with residue M165 forming hydrophobic contacts with S3-B and S2. Finally, E166 is engaged, forming HB and hydrophobic contacts with S3-A and S3-B. Notably, residues S46, M49, L50, and Q189 make contacts with every NIR region (**Figure 3B,C**).

For cluster 5, engagement also starts at the upper loop, with NIR contacting residues E47 and L50 in all successful pathways sampled. This suggests they may serve as early anchor points for NIR binding. Next, binding proceeds towards the lower loop and spans a broader range of residues, with the dominant early interactions in this region concentrated at residues Q189, A191, and Q192. Residues A191 and Q192 form hydrogen bond donating interactions with S3-B and S1-C respectively, while residue Q189 engages S3-A, S3-B and S1-C sites through both HB-donor and HB-acceptor interactions. The ligand is then drawn deeper into the active site by the same residue E166-mediated interactions observed in cluster 4 via S3-A and S3-B, and similar interactions with M49 and M165, followed by interactions between S2 and H41 and between S3-A and S3-B and N142. In cluster 5, residues L50, P168, Q189, and A191 form contacts with every region of NIR (**Figure S8**).

Though cluster 5 spends more time interacting with the lower loop than cluster 4, both pathways proceed from the upper loop to the lower loop before reaching the catalytic loop. Interestingly, the upper loop is the first point of contact even for pathways in cluster 4 which approach from closer to the lower loop. Interactions with S46 M49, M165, or E166, appear to be the next key determinants of binding progression; once contacts with these residues form, the number of total contacts rapidly rises. Contacts with several residues, including S46, E47, L50, H164, and D187 and contact with the S1 subsite of NIR have a low mean contact count (**Figure 3B,C**, **Figure S8**), but are present in over 90% of successful pathways. This suggests that contacts are rapid, or short-lived, yet significant to binding progression. This is supported by prior SAR studies demonstrating that the trifluoroacetamide moiety of S1-A and C and the warhead S3-B region are highly sensitive to modification, with even modest changes (e.g, trifluoro to trichloro) resulting in substantial losses in potency.^69^

Residues M49, M165, E166, and Q189 represent the longest-lived contacts, once formed. Residue P168 forms short-lived contacts in cluster 4, but spends a much longer time in contact, particularly with S2 in cluster 5. The breadth of early-stage contacts captured in the simulations (S1, 24–27, V42, 44–47, 50–51, L141, S144, G170, H172, F181, 186–192, R217) that lead to productive binding relative to the narrow set of interactions in the covalently bound structure 7SI9^66^ (H41, M49, Y54, F140, N142, G143, C145, 163–168, Q189) suggests that many transient encounter complexes are kinetically accessible (**Figure 3**, **Figures S6,S8)**. Taken together, our simulations suggest a dynamic binding mechanism in which upper loop residues E47 and L50 act as early anchor points, followed by lower loop checkpoint contact with Q189, while contacts with the upper loop residues S46, M49, and catalytic residues M165 and E166 contacts initiate settling into the final bound conformation **(Figure S9)**.

### Simulations are consistent with combined conformational selection and induced fit mechanism

Next, we sought to determine whether the binding pathways sample unique Mpro conformations compared to those accessed in the covalently bound or apo states. To extensively explore apo conformations, we employed an exploratory WE protocol, which focused sampling toward pocket conformations which deviate from the initial structure, reaching up to 11 Å RMSD (**Figure S7**). We also conducted a conventional MD simulation of NIR covalently bound to Mpro to characterize the bound state. To visualize the conformational space of these systems, we conducted principal component analysis (PCA) on the Cα coordinates pooled across all three systems. The resulting projection is shown in **Figure 4A**.

**Figure 4:**
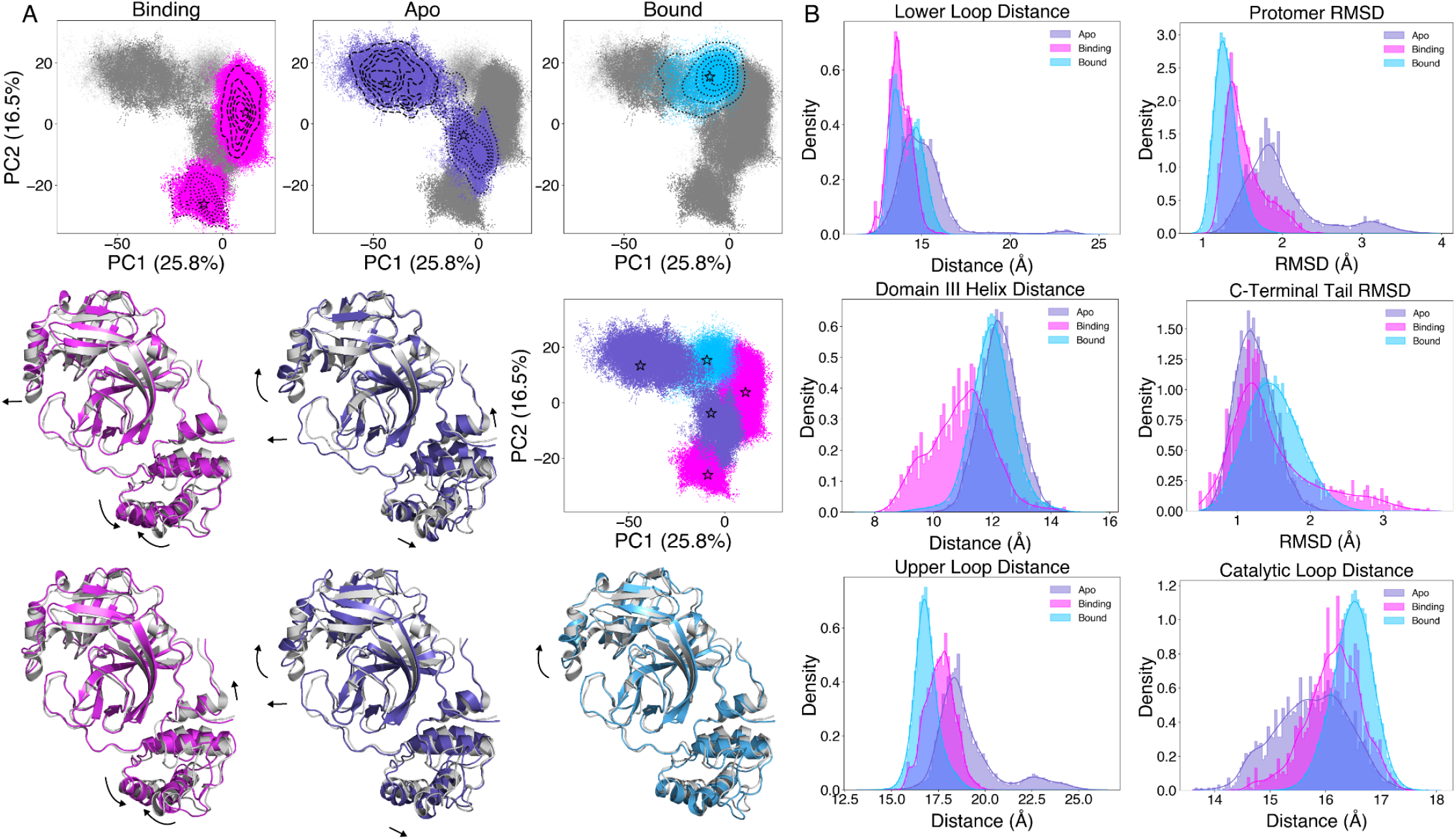
Principal component analysis of binding, apo, and bound simulations. (A) Principal component analysis (PCA) of Cα coordinates projected onto PC1 and PC2. Top row: density contours for the binding (magenta), apo (purple), and bound (cyan) ensembles, each overlaid on the combined conformational ensemble (grey), with the three systems shown combined in the middle-right plot. Stars denote the representative structure of lowest RMSD to each cluster’s mean. Middle and bottom rows: structural superpositions of representative conformations, aligned to the crystal structure PDB 6M03^75^ (grey). The middle-row structure corresponds to dashed line clusters and bottom-row structure corresponds to the dotted line clusters. Black arrows highlight the principal conformational differences relative to the reference. Only a single monomer is shown for clarity. (B) Probability density distributions of six structural measurements across the apo (purple), binding (magenta), and bound (cyan) ensembles: lower loop distance (Cα145–Cα188), protomer RMSD (fit to 6M03^75^), Domain III helix Cα distance (Y237–N274), C-terminal tail (S301-Q306) RMSD, upper loop distance (Cα145–Cα50), and catalytic loop distance (Cα145–Cα168).

While the three systems show overlap in PC space, each also occupies distinct conformational regions. The overlap between the apo ensemble with the NIR binding and bound ensembles is consistent with a conformational selection model, in which bound state-like conformations are accessible within the apo ensemble’s native fluctuations and can be stabilized by a ligand.^5^ Simultaneously, the unique space sampled during the binding ensemble supports the presence of induced-fit conformations, where ligand binding drives conformational rearrangements that would not otherwise occur.

The major differences across all 3 systems are most clearly seen in 5 regions: the upper, lower, and catalytic loops which surround the binding pocket, helices A (L227-Y237) and B (A260-N274) which lie in domain 3 of the protein, and the C terminal tail (S301-Q306) (**Figure 4A,B**). The binding pocket loops, and overall protomer RMSD compared to the crystal structure show most deviation in the apo simulation (**Figure 4A,B**), as expected since the simulation was designed to focus on maximally exploring conformational space. The distance between the catalytic residue C145 to the alpha carbons, L50 (upper loop) and R188 (lower loop) and P168 (catalytic loop) in the apo systems show broad distributions, reaching up to 25 Å in the lower and upper loops, compared to maxima under 20 Å in the bound and binding simulations (**Figure 4B**). Rather than increased maxima in the catalytic loop distance, the broad distribution of the apo system spans decreased distances compared to the binding and bound systems, reaching below 14 Å.

The bound simulation maintained the lowest RMSD compared to the crystal structure (**Figure 4B**) large deviations were not expected for this state, given the constraints of the covalent bond. The bound simulation also maintained the smallest distance between the upper loop and catalytic residue, yet the catalytic loop sampled the largest distance during this simulation spanning 15-18 Å. The binding simulation sampled intermediate distances lower than 15 Å, and greater than 17.5 Å, while the apo system sampled the smaller distances ranging below 14 Å up to 17.5 Å. The lower loop distance range in the bound simulation was smaller than the apo distance, but similar to the binding simulation (**Figure 4B**).

One unique motion that was observed only in the binding simulations involved the A and B helices of domain III, which seem to come closer together compared to the crystal structure (**Figure 4A**). This is in contrast to the apo system, where these helices appear to move in unison (**Figure 4A**). The distribution of distances between Y237 and N274, the tips of domain III’s helix A and helix B, respectively, reveal a left skew, where the binding simulation spans a greater range, from 8 Å to over 14 Å, compared to the bound and apo which do not reach below 9 Å or 10 Å, respectively. This distal site has been proposed as a putative allosteric site.^70–73^ The C terminal tail is another region showing larger deviation in the binding simulations compared to the apo or bound states (**Figure 4B**). This phenomena has been observed in crystal structures, where those with only a single protomer binding site occupied showed a large perturbation.^74^ Both of these regions are far from the binding pocket and demonstrate that although our WE progress coordinate was focused on distance and RMSD of the ligand alone, we are able to capture these long-range allosteric motions involved in the binding process.

Collectively, the overlapping conformations between apo, binding, and bound states suggest some conformational selection is present in the apo state, whereas the unique regions of conformational space only detected in the binding simulations are consistent with induced fit mechanisms also playing a role. The distances and RMSDs measured generally follow a trend in which binding sits between the apo and bound distributions, with the exception of the binding pathway-specific dynamics observed between helices in domain III and the C-terminal tail.

### Simulation and Experiment Establish a Cohesive Model of Binding Kinetics

Next, we sought to integrate the results from WE simulations and NMR titrations for a comprehensive characterization of the NIR-Mpro binding pathway and dynamics. Several residues showed direct agreement between experiment and simulation. For example, residues V42, T45, S46, 142-144, H172, V186, T190, and Q192 are contacted in the simulations and exhibit intermediate exchange in the WT NMR titration (**Figure 5A**). Residue H163, also contacted in the simulations, is unassigned in the WT spectra, but exhibits intermediate exchange in the catalytically inactive NMR titration. Residue G170 exhibits slow exchange in the NMR, is present in 100% of pathways in cluster 5, but only occurs in one pathway in cluster 4. This residue sits on the lower loop, and based on the initial NIR position for cluster 5 pathways, NIR must pass over G170 to reach the binding pocket opening (**Figure 2A**). NMR chemical shifts represent population-weighted ensemble averages, whereas the distinct pathways captured in the simulations help disentangle individual conformational states and pathways.

**Figure 5:**
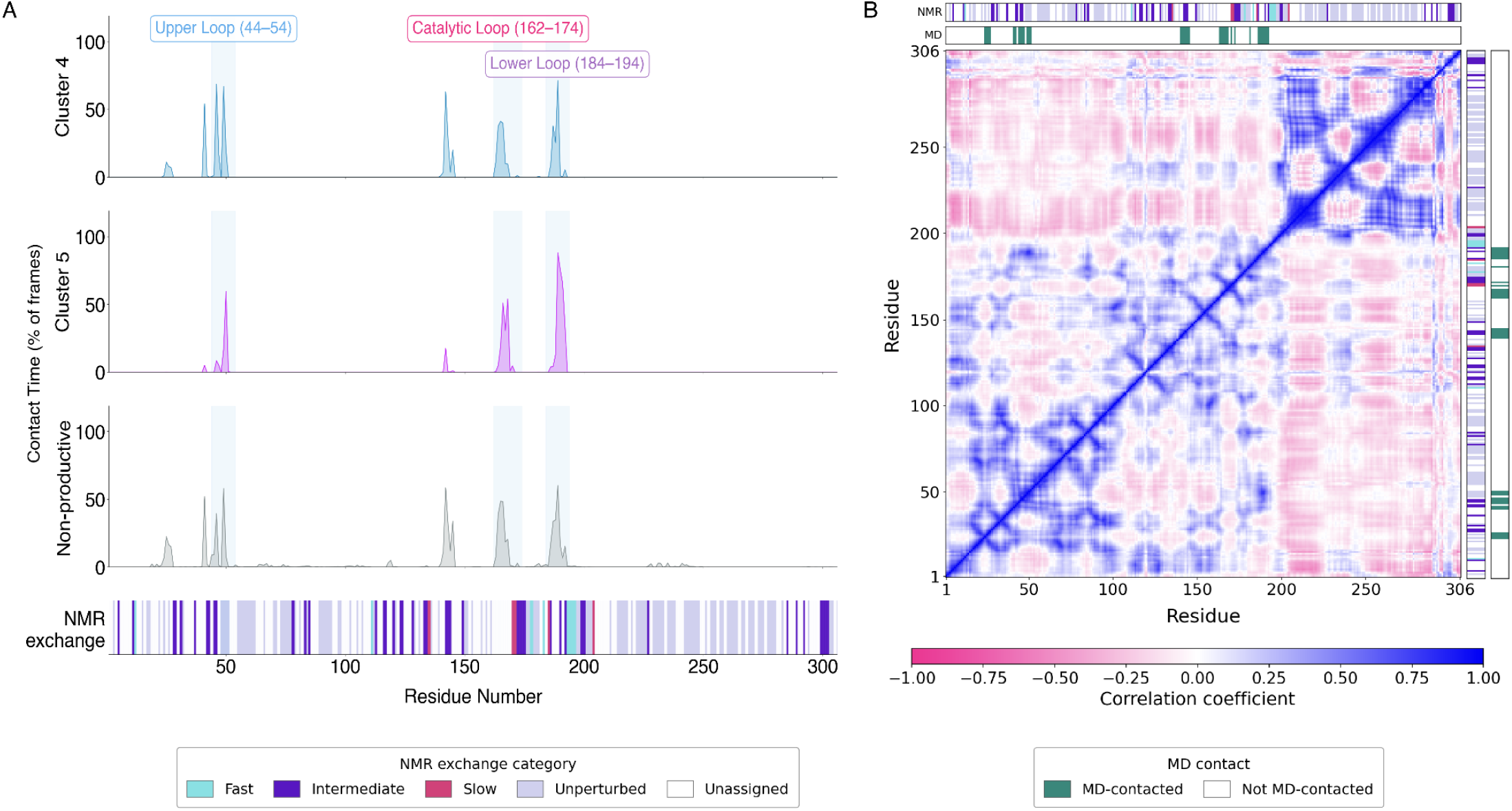
Comparison of residue-level contacts and dynamics during binding simulations and NMR titrations. (A) Contact time between protein residues and NIR, calculated as the percentage of trajectory frames in which a contact is present, are shown for cluster 4 (magenta) and cluster 5 (cyan) and non-productive (grey). Shaded regions denote key structural elements including the upper, catalytic and lower loop. Bottom plot shows NMR exchange classification for each residue. (B) Residue–residue correlation matrix from binding simulations showing 306 residues of the protomer NIR binds to. Top and right-side margin show the NMR exchange category (colored as in A) and whether the residue was contacted by NIR in the MD simulations.

Some of the residues contacted in the MD simulations were unperturbed in the NMR. Residues T24, T26, M49, L50, N51, and F181 were unperturbed in the WT, while M49 exhibited fast exchange in the C145A variant. Residues T24, T26, L50, and F181 were unassigned in the C145A and N51 was similarly unperturbed. Of these, L50 was found to be a key early anchor point forming a transient contact in the simulations. L50 lines the pocket opening, and L50F was found to be a hyperactivating variant,^68^ which can compensate for activity loss induced by resistance mutations such as E166V. Therefore, participating in the early stages of substrate binding offers a plausible explanation, as it does not directly interact in the bound state. The ^1^H-^15^N HSQC assignments are limited to the nitrogen atoms of the residue backbone, therefore it is possible the leucine side chain interactions with NIR may not impact the backbone chemical environment. Residue M49 exhibits this phenomenon as well, forming hydrophobic interactions during simulations which may not affect the residue backbone. Furthermore, MD timesteps capture motions occurring as fast as the femtosecond timescale, therefore some transient contacts detected in the simulations may be too fast to be observed in NMR.

The residues perturbed in NMR but not observed to make contact in the simulations, including all of the fast exchange residues and some of the intermediate and slow exchange residues, are greater than 5 Å from the bound state NIR in the crystal structure. These may represent non-specific direct interactions in solution, or residues experiencing a new chemical environment in response to conformational changes during binding. Non-specific interactions are mostly unaccounted for in our simulations based on the nature of the WE approach, which focuses sampling along a progress coordinate. Many pathways that do not directly approach the target state are terminated (and their weight re-assigned) to focus computational resources toward sampling the pathway of interest. This approach enables us to sample events that occur on otherwise inaccessible timescales.

Some of the non-contacted yet NMR perturbed residues can be directly attributed to allosteric motions observed in the simulations. For example, the slow exchange residue V204 sits in the domain III helix spanning residues I199-N214, and is not solvent accessible, therefore direct binding interactions are unlikely to be the source of this perturbation. This helix is orthogonal to, and highly correlated to, the helix B A260-N274 found to be compressed closer to helix A L227-Y237 during our binding simulations (**Figure 5B**, **Figure S10-11)**.

Residue V204 is also highly correlated to the loop connecting the last helix before the C-terminal tail, including residues G275-F291, and anticorrelated to several residues spanning the N-terminal domain (**Figure 5B**, **Figure S10-11)**. The other slow exchanging residues including T135, G170, V171, and F185 are near the bottom of the binding pocket, with G170 and V171 being part of the catalytic loop and F185 part of the lower loop. While only G170 was observed to directly interact with NIR in cluster 5, these residues are within a 5 Å radius, and there are strong correlations throughout this region particularly between T135 and the lower loop residues M165-G174, catalytic loop residues F181-F185, and the loop connecting the catalytic domain to domain III residues A193-T199 (**Figure 5B**, **Figure S10-11**), providing a plausible explanation for their chemical shift perturbation other than direct interaction. Individually, WE-MD and NMR CSP provide complementary but incomplete views of the binding process, and though they cannot definitively resolve every complexity of the drug association ensemble, combining information from each yields a more complete mechanistic understanding.

## Conclusion

In this study, we resolve elements of the kinetically complex mechanism of molecular recognition via NIR binding to SARS-CoV-2 Mpro. Solution NMR titrations combined with WE enhanced sampling simulations offer a synergistic perspective of the dynamic process. Our ^1^H-^15^N HSQC titrations demonstrate that ligand binding involves a heterogeneous mixture of fast, intermediate, and slow exchange regimes in residues distributed across the active site and distal regions of the protein. The WE-MD simulations generated 563 continuous binding pathways originating from two unique initial ligand placements, producing two dominant, statistically independent routes. Our simulations reveal a transient mechanism in which residues E47 and L50 in the upper loop of the Mpro binding site serve as early anchor points for NIR, which are followed by interactions with Q189 in the lower loop, positioning NIR for contact with S46, M49, M165, or E166 which lead to rapid rearrangement into the final bound pose through a combined mechanism involving conformational selection followed by induced fit. Eleven residues contacted in MD showed intermediate or slow exchange during NMR titrations, including the key residue S46. The agreement between these experimental observations and prior literature provides support for the simulations and their mechanistic interpretation.

Both NMR and WE provide distinct, yet complementary insights into the binding mechanism. The HSQCs provide an experimentally grounded measurement indicating the chemical environment changes experienced by each residue during binding. The simulations reveal three dimensional explanations for many of the observed perturbations. In this large system, where several peaks cannot be assigned due to spectral crowding, MD fills in the missing information about those residues. The HSQC spectra show the breadth of interactions occurring with regions of the protein not detected by the pathways sampled in MD restricted to a predefined progress coordinate. The simulations reveal allosteric motions correlated with residues experiencing slow exchange in NMR, particularly residue V204, which is far from the binding site and not solvent exposed, and therefore likely cannot directly interact with NIR. The perturbations detected with NMR represent a population-weighted average of the true ensemble, while the simulations offer analysis of individual atomistic pathways. This comprehensive characterization of binding pathways sheds light on the structure-kinetic relationship between NIR and Mpro, providing new potential avenues for next-generation inhibitor design beyond optimization of just the final bound conformation. Though NMR and MD have long been viewed as synergistic approaches,^16–19^ combining them in this way to characterize drug binding pathways has not been explored. Future developments tuning simulation parameters with experiments, and expanded NMR analysis, will likely provide even deeper mechanistic insights.

## Methods

### Experimental Methods

#### Plasmids

The plasmid encoding WT Mpro was a gift from Dr. Ashootosh Tripathi (University of Michigan) which contains the WT Mpro sequence cloned into a pET-His_6_-SUMO vector. The C145A mutant was generated from the WT construct by side directed mutagenesis (SDM) with forward and reverse primers GAACGGCTCAGCGGGCTCTGTAG and AAAAAAGAGCCTTTGATAGTG respectively (Annealing temperature 58 °C) purchased from New England Biolabs (NEB) using an SDM kit from NEB.

#### Protein Expression and Purification

Expression and purification of ^15^N labeled protein was carried out using previously established protocols.^76^

Briefly, the plasmid encoding Mpro WT or C145A mutant was transformed into *E. coli* BL21(DE3). The cells were grown in M9 minimal medium consisting of 6 g/L Na2HPO4, 3 g/L KH2PO4, 0.5 g/L NaCl, 10 g/L MgSO4 0.6 g/L CaCl2, 0.4 g/L thiamine, 0.12 g/L FeS04, 4 g/L glucose and 1 g/L ^15^NH_4_Cl with 1 ml of 1000x ampicillin (100 mg/ml) to a final working concentration of 100 µg/ml at 37 °C to an OD600 of 0.8-0.9 and cooled down to 16 °C for about 30 mins at which point OD600 reached around 1.0. Protein expression was then induced by addition of 1 mM isopropyl-B-D-galactopyranoside (IPTG) for approximately 18 hours at 16 °C. Cells were harvested at 3,500 rpm for 40 mins and lysed by sonication in buffer A consisting of 50 mM sodium phosphate buffer at pH 7.4, 250 mM NaCl and 10% glycerol. The lysate was pelleted by centrifugation at 20,000 rpm for 40 mins and the supernatant was incubated with nickel resin for 1 hour at 4 °C then passed through a gravity column (15 ml) to pack the resin and the flow through was collected. The resin was washed with buffer A containing 25 mM of imidazole for 7 column volumes. After washing, the protein was eluted with 2 column volumes of buffer A containing 250 mM imidazole. The eluted protein was dialyzed in 50 mM Phosphate at pH 7.4, 150 mM Sodium Chloride, 1.5 mM Sodium Azide, 1 mM TCEP, 10% glycerol and SUMO protease overnight. After dialysis the protein was incubated with nickel resin to capture the SUMO-his6 tag while untagged protein was collected from the flow through. Protein was concentrated to 0.5 mL volume for size exclusion chromatography using a Superdex 75 increase 10/300 GL (Cytiva) column equilibrated with NMR buffer 10 mM Sodium Phosphate at pH 7.0, 0.5 mM TCEP. The fractions containing Mpro were pooled and concentrated. The A280 of the protein was obtained using a nanodrop and the final concentration of Mpro was determined using an extinction coefficient of 65,780 M^-1^cm^-1^.

#### NMR methods

NMR samples were prepared by addition of 3% D_2_O to protein at concentration of 300 µM in NMR buffer consisting of 10 mM sodium phosphate at pH 7.0, 0.5 mM TCEP to a final volume of 500 µl for WT and 180 µl for C145A in a 5 mm or 3 mm NMR tubes, for WT and C145A respectively. Spectra were acquired for increasing concentrations of NIR (Sigma) ranging from 0 - 1.25x for WT and 0 - 3x for C145A on an 800 MHz Bruker ADVANCE NEO spectrometer at 25 °C for WT and 37 °C for C145A using a standard TROSY (Transfer Relaxation Optimized Spectroscopy) HSQC pulse sequence (trosyetf3gpsi). NMR spectra were processed with NMRFx Analyst^77^ and analyzed with Sparky^78^. Backbone resonances were assigned using BMRB, IDs 50780, 51455 as references.^47,52^

### Computational Methods

#### System Preparation

All unbound starting states were modeled utilizing the crystallographic structure PDB ID: 6M03^75^, the covalent bound state was PDB ID: 7SI9.^66^ Protons were added at pH 7.4 with the H++ web server.^79^ For non-covalent simulations, NIR was modeled with RDKit from the following SMILES string: CC1(C@@H]2[C@H|1[C@H](N(C2)C(=0)[C@H](C(C)(C)C)NC(=O)C(F)(F)F)C(=O)N[C@@H](C|C@@H|3CCNC3=0)C#N)C.

For the covalent simulation, NIR is connected to C145. Therefore, prior to charge calculation, the cysteine residue was capped with acetyl (ACE) and N-methylamide (NME) groups to reproduce the local chemical environment of the attachment site. To determine an accurate electrostatic representation, the ligand was first optimized at the HF/6-31G* level of theory using Gaussian 16.^80^ The resulting electrostatic potential (ESP) was then used to derive restrained electrostatic potential (RESP) atomic partial charges and assign atom types using the Antechamber and Prepgen tools in Amber24.^81^ NIR was subsequently parameterized using the General Amber Force Field 2 (GAFF2).^82^ Remaining bonded parameters identified and generated using parmchk2. The resulting parameters were incorporated into a force field modification file used for the final system assembly.

Next, the protein and ligand were integrated into a single system and tleap was used for solvation and to generate topology and coordinate files. The ff14SB^83^ force field was applied to the protein atoms. Each system was solvated in a truncated octahedral box of TIP3P^84^ water molecules, extending 20 Å from the protein. The systems were neutralized and adjusted to physiological ionic concentration of 150 mM with Na+ and Cl− ions.

Each of the systems were subjected to the following steps with pmemd.cuda.^85,86^ First, a two step minimization was carried out. In the first step, we minimized the bulk solvents and ions using 10,000 minimization cycles with a harmonic positional restraint of 500 kcal/mol·Å^2^ applied to all protein and ligand heavy atoms. In the second step, a fully unrestrained minimization was performed over 100,000 cycles.

Following minimization, the system was heated in 7 sequential stages of 50 ps each (350 ps total), raising the temperature from 0 K to 310 K. The first heating stage used a timestep of 1 fs, while subsequent stages used a timestep of 2 fs. Incremental heating was carried out in the NVT ensemble using a Langevin thermostat with a collision frequency of 1 ps^-1^. Following heating, the system was equilibrated for 1 ns in the NPT ensemble under constant pressure conditions (1 atm) using a Monte Carlo barostat, with no positional restraints applied. Production steps were carried out using the same protocol as equilibration steps. The covalent simulation was conducted for 100 ns. For binding simulations, the final configuration from equilibration served as the initial state for WE simulations, where the same production protocol was used with 100 ps resampling intervals. All input files used for the minimization, heating, and equilibration steps are available in the accompanying GitHub repository.

#### Weighted Ensemble Simulation

Weighted ensemble (WE) simulations of NIR binding were carried out under non-equilibrium steady-state conditions using the open-source WESTPA^87^ software package. Under steady-state conditions, upon reaching the target state, the walker is terminated and a new walker is initialized from a new basis state with identical statistical weight.^62^ Progress was measured via two coordinates. 1) The distance between the sulfur atom of the Mpro catalytic cysteine 145 and the nitrogen atom of the NIR nitrile warhead 2) RMSD of the ligand after alignment of the protein to PDB ID: 7SI9.^66^ Progress coordinates were calculated using CPPTRAJ.^88^ The target state for recycling was defined as both progress coordinates reaching ≤ 5 Å. The progress coordinate space was discretized using the following two-dimensional binning scheme: RMSD bins [0, 1.0, 2.0, 3.0, 5.0, 5.5, 6.0, 6.5, 7.0, 8.0, 9.0, 10.0, 15, 20, 25, 30, 35, 40, infinity] Å and distance bins of [0, 4.0, 5.0, 6.0, 7.0, 8.0, 10.0, 15, 20, 25, 30, 35, 40, infinity] Å. The simulation was initialized from 64 basis states with unique ligand positions. To generate ligand positions, the protein’s position remained fixed while the ligand was projected onto a random point on a sphere with a radius of 20 Å centered on the protein’s center of mass. A random rotation was also applied to the ligand according to the Fast Random Rotation algorithm described by J. Arvo.^89^ Each of these were prepared with the minimization, heating, and equilibration procedures described above. Basis states were each assigned equal statistical weight. WE resampling was performed at fixed time intervals of 100 ps, maintaining a target of 8 walkers per bin.

For exploratory simulations of the apo Mpro system with no ligand, steady-state conditions were not applied. A single equilibrated starting state was used. A two-dimensional progress coordinate consisted of: 1) Full backbone RMSD, and 2) RMSD of just the pocket residues including 26–28, 39, 41, 49, 54, 140, 141–144, 146–147, 163–168, 172, 181, and 186–192. All RMSD values were computed against the first frame of the starting state. To maximize the exploration of novel states, the MABBinMapper (Minimal Adaptive Binning)^90^ scheme was employed within a bin boundary of [0,infinity]. The adaptive mapper was configured to maintain eight bins along each dimension of the progress coordinate, prioritizing the sampling of walkers that exhibited increasing RMSD values.

### Simulation Analysis

#### Simulation Convergence

Simulation convergence was assessed using the WESTPA w_ipa analysis tool to calculate the target-state flux and approximate the rate constant. For this analysis, bound and unbound states were defined to match the target state bins used to drive the WE simulation: a two-dimensional RectilinearBinMapper with boundaries [0.0, 5.0, inf] on each progress coordinate dimension was used to classify walkers as bound coordinates, < 5 Å, or unbound coordinates, > 5 Å. Kinetics were evaluated with a cumulative evolution scheme, giving a rolling estimate of the target flux and rate constant, with 95% confidence intervals, at each iteration. Simulations were continued until the estimated rate constant reached convergence, as indicated by the absence of significant fluctuations over subsequent iterations.

#### Pathway Selection

LPath^91^ match and extract steps were used to identify successful trajectories. For the binding simulations, successful transitions were identified using a discretized state scheme of [0.0, 5.0, ∞] Å for both progress coordinates. This reports all trajectories that reached the target state, including both recycled (terminated) walkers and those that entered the target state but were not immediately recycled due to leaving the target state before the end of the resampling interval. For the exploratory apo simulations, successful "events" were defined post-hoc using a probability-based cutoff derived from the population histograms. Which resulted in the following scheme: [0.0, 3, ∞], and [0.0, 4, ∞] for backbone and pocket rmsd, respectively.

Complete and continuous trajectories were reconstructed by tracing successful walkers back to their respective initial states.

To categorize the diverse binding mechanisms observed, hierarchical clustering was performed on the successful binding trajectories using a lineage-weighted Jaccard distance, calculated from the sequence of walker IDs visited by each trajectory and weighted by 1/iteration to emphasize early-stage divergence trajectories. This form of weighting was chosen over an unweighted scheme and one that instead grew more heavily weighted toward later iterations, based on cluster separation measured by silhouette score; the early-stage (1/iteration) weighting gave the best score. Applying this distance metric resulted in 7 clusters, two clusters 4 and 5, together accounted for ∼90% of the total recycled trajectory weight and were carried forward as representative pathways for detailed analysis. Full clustering code is available in the accompanying GitHub repository.

#### Trajectory Analysis

Protein-ligand interaction analysis was conducted using the ProLIF (Protein-Ligand Interaction Fingerprints) Python package^65^. Contacts were counted as the number of frames each residue spent within the cutoff distance of the ligand per residue. The interaction fingerprints were configured to specifically detect hydrogen bond donors and acceptors, hydrophobic contacts, pi-stacking, cationic and anionic interactions, and van der Waals contacts. ProLIF classifies a hydrogen bond as a polar heavy atom (N, O, S, or protonated N) bearing a donatable hydrogen positioned within 3.5 Å of, and oriented toward, a suitable acceptor atom (N, O, or F), with the donor-hydrogen-acceptor angle constrained to 130-180°. Hydrophobic contacts are identified by proximity, within 4.5 Å, between non-polar aromatic and aliphatic carbons, divalent sulfurs, and the halogens Br and I, excluding any carbon directly bonded to nitrogen, oxygen, or fluorine. Van der Waals contacts are characterized by an atom pair separated by an interatomic distance no greater than the sum of their van der Waals radii, using ProLIF’s default MDAnalysis-derived radii values.^92,93^

Distance, RMSD and PCA were calculated using CPPTRAJ.^88^ For RMSD calculations, the backbone atoms corresponding to the protomer with which the ligand interacts were first aligned to the reference structure PDB 6M03^75^, and the resulting RMSD of the Cα atoms was calculated using the nofit option.

#### Principal Component Analysis

Principal component analysis was performed on Cα coordinates. Using CPPTRAJ^88^, a coordinate matrix of Cα atoms across all frames aggregated from apo, binding, and bound simulations was constructed. The mean structure was computed through iterative alignment of the coordinate matrix to the averaged structure followed by re-averaging and re-fitting until the Cα RMSD between successive iterations fell below 1 × 10^-3^ Å. The covariance matrix was computed and diagonalized using numpy.^94^ Representative conformations were identified by clustering the trajectory in PCA space using a Gaussian mixture model (GMM).^95^ Within each cluster, the mean structure was calculated and the structure with the lowest RMSD to the cluster mean was selected as the representative conformation and used for visualization.

### Data availability

Additional data supporting the findings of this study are presented in the SI Appendix. Input files and scripts used to set up and run the molecular dynamics simulations are available on GitHub at https://github.com/SztainLab/NIR-mpro-binding

## Acknowledgements

We thank the Advanced Cyberinfrastructure Coordination Ecosystem: Services & Support (ACCESS) program, which is supported by U.S. National Science Foundation grants #2138259, #2138286, #2138307, #2137603, and #2138296. Portions of this work used the Expanse resource at the San Diego Supercomputer Center https://doi.org/10.1145/3437359.3465588 supported by ACCESS allocation BIO240047. We thank Dr.

Ashootosh Tripathi for providing the plasmid for Mpro. We also thank Jeff Thompson for providing code that was adapted for generation of Figure S5.

## Author Contributions

T.S. designed research; D.S.P., G.A., S.M., and S.H.L. performed research; D.S.P., G.A., Y.F., S.M., S.H.L., and T.S. analyzed data; and D.S.P., G.A., and T.S. wrote the paper.

## Competing Interests

The authors declare no competing interests.

## Notes

### Competing Interest Statement

The authors have declared no competing interest.

